# Automated remote focusing, drift correction, and photostimulation to evaluate structural plasticity in dendritic spines

**DOI:** 10.1101/083006

**Authors:** Michael S Smirnov, Paul R Evans, Tavita R Garrett, Long Yan, Ryohei Yasuda

## Abstract

Long-term structural plasticity of dendritic spines plays a key role in synaptic plasticity, the cellular basis for learning and memory. The biochemical step is mediated by a complex network of signaling proteins in spines. Two-photon imaging techniques combined with two-photon glutamate uncaging allows researchers to induce and quantify structural plasticity in single dendritic spines. However, this method is laborious and slow, making it unsuitable for high throughput screening of factors necessary for structural plasticity. Here we introduce a MATLAB-based module built for Scanimage to automatically track, image, and stimulate multiple dendritic spines. We implemented an electrically tunable lens in combination with a drift correction algorithm to rapidly and continuously track targeted spines and correct sample movements. With a straightforward user interface to design custom multi-position experiments, we were able to adequately image and produce targeted plasticity in multiple dendritic spines using glutamate uncaging. Our methods are inexpensive, open source, and provides up to a five-fold increase in throughput for quantifying structural plasticity of dendritic spines.

## Introduction

Structural changes in dendritic spines, tiny postsynaptic protrusions on the dendritic surface of neurons, are considered to be the basis of synaptic plasticity, learning, and memory [1–5], Among several forms of spine structural plasticity, structural long-term potentiation (sLTP) of single dendritic spines has been extensively examined as a structural correlate of functional LTP (fLTP), an electrophysiological model of learning and memory [3, 5, 6], Applying two-photon glutamate uncaging at a single dendritic spine induces a rapid and long lasting spine enlargement at the stimulated spine, but not the surrounding spines [3], The signaling cascades necessary for sLTP have been studied by combining sLTP imaging with pharmacological and genetic manipulation. Both sLTP and fLTP depend on NMDA-type glutamate receptors (NDMAR), Ca2+/Calmodulin-dependent kinase II (CaMKII) and several small GTPase proteins including Ras, Cdc42, Rac1 and RhoA [4, 7-11], However, due to the low-throughput nature of the measurement, the study of sLTP has been limited to only a few proteins, and among more than 1000 proteins expressed in spines [12-14], it is largely unknown which ones are necessary for spine structural plasticity.

The quantification of long-term structural plasticity of dendritic spines requires imaging single spines over extended periods of time (typically ∼1 h), and it is necessary to continuously refocus to the target spines. Moreover, in general, imaging and stimulating multiple dendrites over long periods of time has been difficult with regular two-photon microscopy, limiting the throughput of the quantification of spine structural plasticity. Thus, it was necessary to develop a system that allows for 1) automatic focus and drift correction for long-term tracking of dendritic spines and 2) imaging and stimulation of several regions of interests (ROIs).

Although automated focusing for microscopy is a well-studied topic in literature [15-22], most algorithms are designed for highly specific imaging modes and preparations. These techniques have been tested under well-defined parameters, therefore, their application in a novel paradigm often results in an overwhelming amount of trial-and-error. Algorithms tend to be uniquely suited to either bright-field, phase, or fluorescence microscopy [23], A lack of functional specificity for software focusing has led some groups to development more inclusive algorithms [20], but the wide variety of imaging setups and biological preparations leaves these attempts incomplete. Hardware focusing systems that correct focus by tracking coverslip location are also commercially available [24], but are incapable of correcting focus drift due to sample deformation which occurs in soft neuronal tissue such as brain slices. In order to optimize software focusing for dendritic spine imaging, an automated algorithm selection tool is necessary to best adapt to the optical parameters and tissue characteristics in the experiment. In addition, since live neuronal tissue can be highly sensitive to objective movements, it is necessary to use a focusing system without objective movements, e.g. an electro-tunable lens (ETL) [25–27],Based on the open-source MATLAB imaging suite Scanimage [28] and an ETL, we have developed an automated system to stimulate and image several individual spines over extended periods of time. With the implementation of an ETL and custom tracking software, our system avoids any artifacts caused by objective or stage movement. We demonstrate that this system allows us to rapidly quantify sLTP in large number of spines.

## Results

In order to achieve rapid and reliable focusing required for auto-focusing system, we employed ETL to our two-photon microscope by placing it at a conjugate plane of the back-focal plane of the objective (Fig 1A). A pre-ETL lens resizes the beam to fit the full aperture of the ETL, while two more lenses serve to resize the beam to fit the galvanometers. In this setup, regardless of ETL shape, the beam size is constant at the galvanometers and the back aperture plane of the objective. This setup minimizes the loss of beam intensity and spherical aberration.

**Fig 1.**
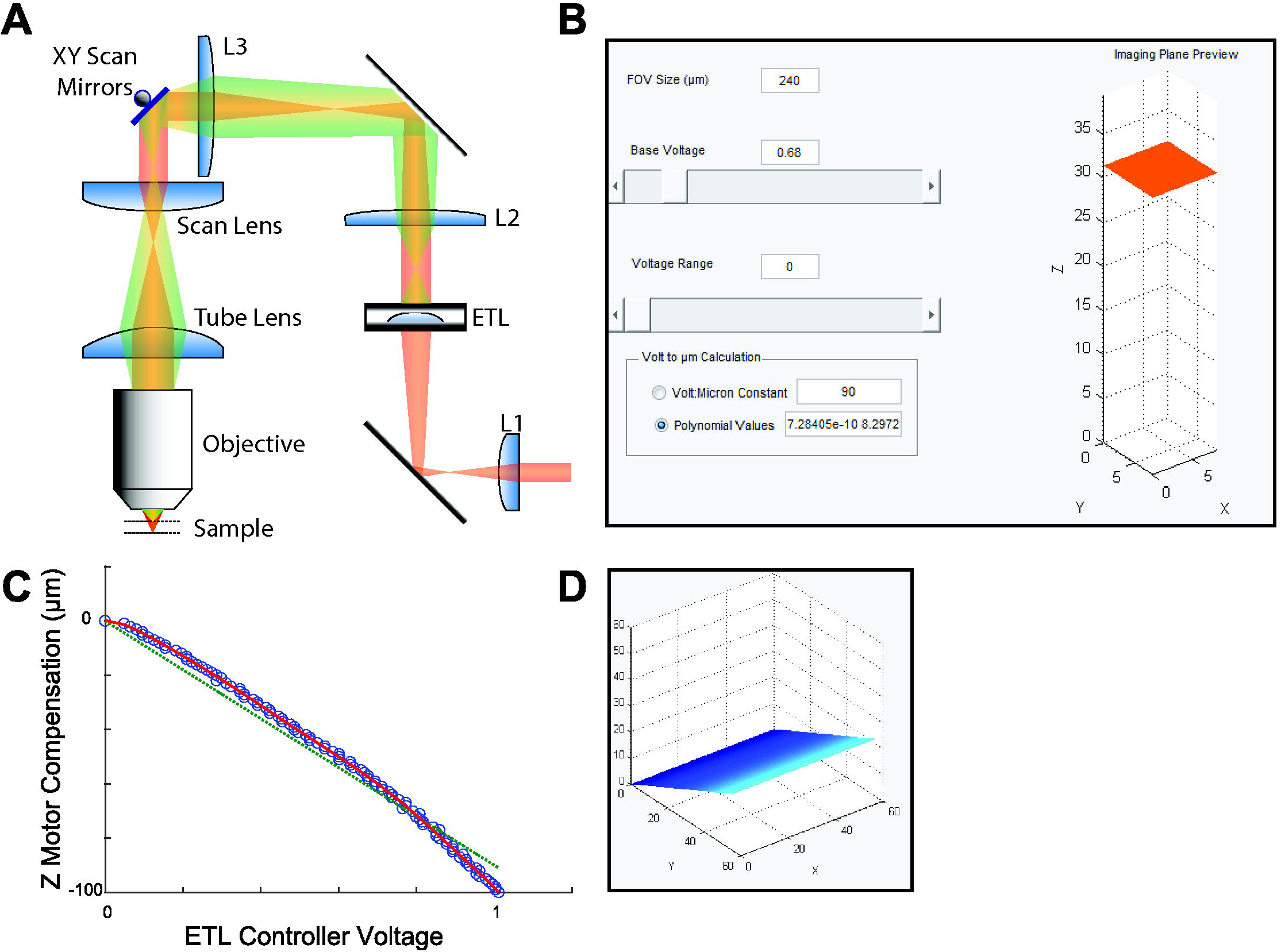
ETL installed in the excitation path and controlled via software GUIs. (A) An ETL in the excitation pathway shapes the incoming beam for remote focusing. Beam shape (red/green) is controlled by curvature of the ETL. L2 and L3 lenses are placed to conjugate the ETL to the back aperture of the objective lens. (B) A GUI controls ETL shape. Voltage values are translated to Z position, and the relative shift of the imaging plane is previewed in a 3D graphic. Z values corresponding to voltages are set either by a linear constant or polynomial curve. (C) ETL voltages are correlated to linear stage movements using an automated alignment routine. (D) Tilted imaging is previewed by adjusting voltage range, as the ETL voltage is altered in phase with the slow-scanning galvanometer.

We controlled the focal length of ETL by changing current passing through the lens using a custom interface built into Scanimage using Matlab (Fig 1B). Initial tuning of the ETL is accomplished by stepwise movement of a Z motor combined with an autofocusing step using solely the ETL. The resulting voltages used to control the ETL are automatically associated with Z displacement values, while a conversion function is set using either a linear constant or polynomial fitted curve (Fig 1B,C). A 3D preview of the imaging plane shows the relative position of the Z plane as a result of ETL offset. Finally, by varying the ETL current in phase with the galvanometer scanning cycle, we were able to tilt the imaging plane, thus allowing users to capture objects which would otherwise require multiple Z slices [27] (Fig 1D).

In order to make long-term imaging possible, we added an interface to compensate for both axial and lateral drift (Fig 2B). Before the start of imaging, users define the range, step size, and frequency parameters for an autofocus routine. When Z-stacks are collected, images are automatically used for auto-focusing. A region of interest (ROI) is defined around the dendritic spine to be stimulated. In order to expand the applicability of our autofocus module to various imaging setups, we have included 30 different autofocusing algorithms, previously described by Pertuz et al. [29], To test the appropriate algorithm for our purpose, we designed an additional application which tests each algorithm against a set of pre-acquired Z stack images, comparing both accuracy and computation time (Fig 2A). In order to ensure that the autofocus algorithm is running normally and using the appropriate part of an image, each collected slice is displayed along with its respective ROI and Z position (Fig 2D). Finally, lateral drift is measured by comparing the imaged position with a reference image by calculating cross-correlation (8), and corrected by immediately shifting the galvanometer scanning angle. While both autofocus and lateral drift correction speeds are dependent on the pixel count of an image, the speed of calculation for a 128x128 pixel image is consistently less than 10 ms (Fig 2D), far exceeding the speed required for most imaging setups.

**Fig 2.**
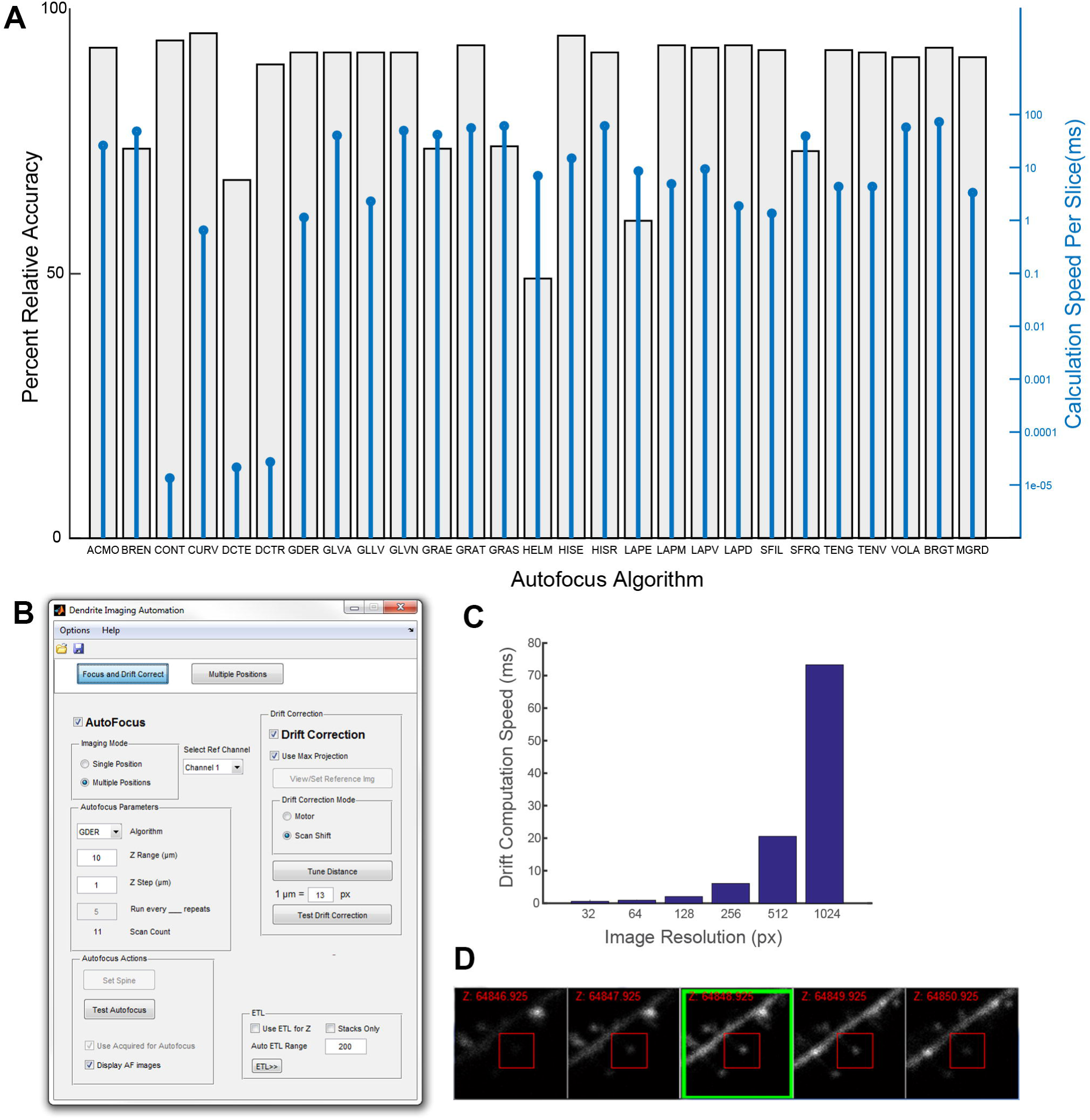
Custom autofocus and drift correction parameters track spines overtime. (A) Sample result of the autofocus algorithm selection tool based on 30 Z stacks with 6 slices each. All algorithms adapted from Pertuz, et al. [29], Abbreviations expanded in Table 1. Percent relative accuracy (gray bars) indicates standardized mean distance of Z position selected by algorithm vs target. 100% = no distance, 0% = maximum distance, 50% = distance if position is picked at random. Blue lines indicate average time to calculate focus value for each slice. (B) Focus and drift correction is controlled through a GUI. Users identify an algorithm, Z range and amount of steps, and whether extra images are collected for autofocus. Drift correction can be enabled to use galvanometers (scan shift) or motor repositioning. Users also have the option to enable or disable the ETL. (D) Reference-based drift correction speed is correlated with pixel size. Image resolution indicates pixel count for one dimension of a square image. (C) Live updates inform users of the selected focus position (green box) and spine ROI used to determine relative focus value (red box).

**Table 1.**
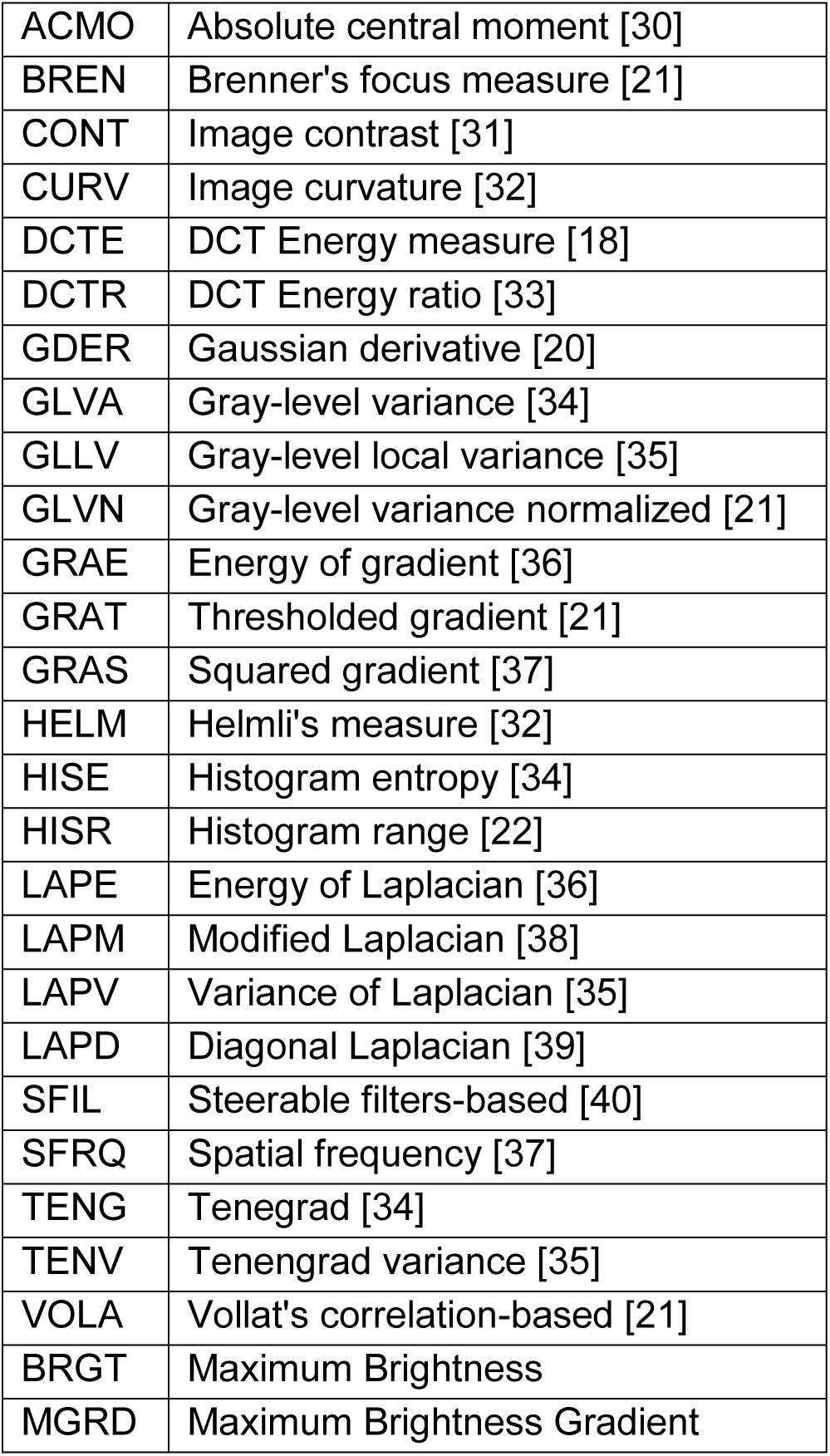
**Abbreviations for autofocus operators**

In addition, we designed a user-friendly interface to rapidly find, store, image, and stimulate multiple positions (Fig 3). Either motor or scanning controls are used to identify dendritic spines for photostimulation (glutamate uncaging). As positions are defined, reference images are automatically collected (Fig 3D). Once all positions are defined, all motor positions are mapped to galvanometer scanning angles (Fig 3A). A separate window allows users to design a timeline for their experiment (Fig 3B). The timeline allows users to control imaging frequency, duration, and define when glutamate uncaging will occur to stimulate dendritic spines. Timeline events can be staggered between positions to avoid conflict between successive uncaging or exclusive imaging events. If the amount of defined positons exceeds the maximum which could be concurrently imaged within a given time constraint, new positions are rotated into the imaging sessions as imaging for other positions is completed (Fig 3B). As imaging proceeds, each position is continuously updated using its reference image (Fig 3D), while an automatic re-alignment of the photoactivation ROI to the cell membrane immediately precedes photoactivation (Fig 3E).

**Fig 3.**
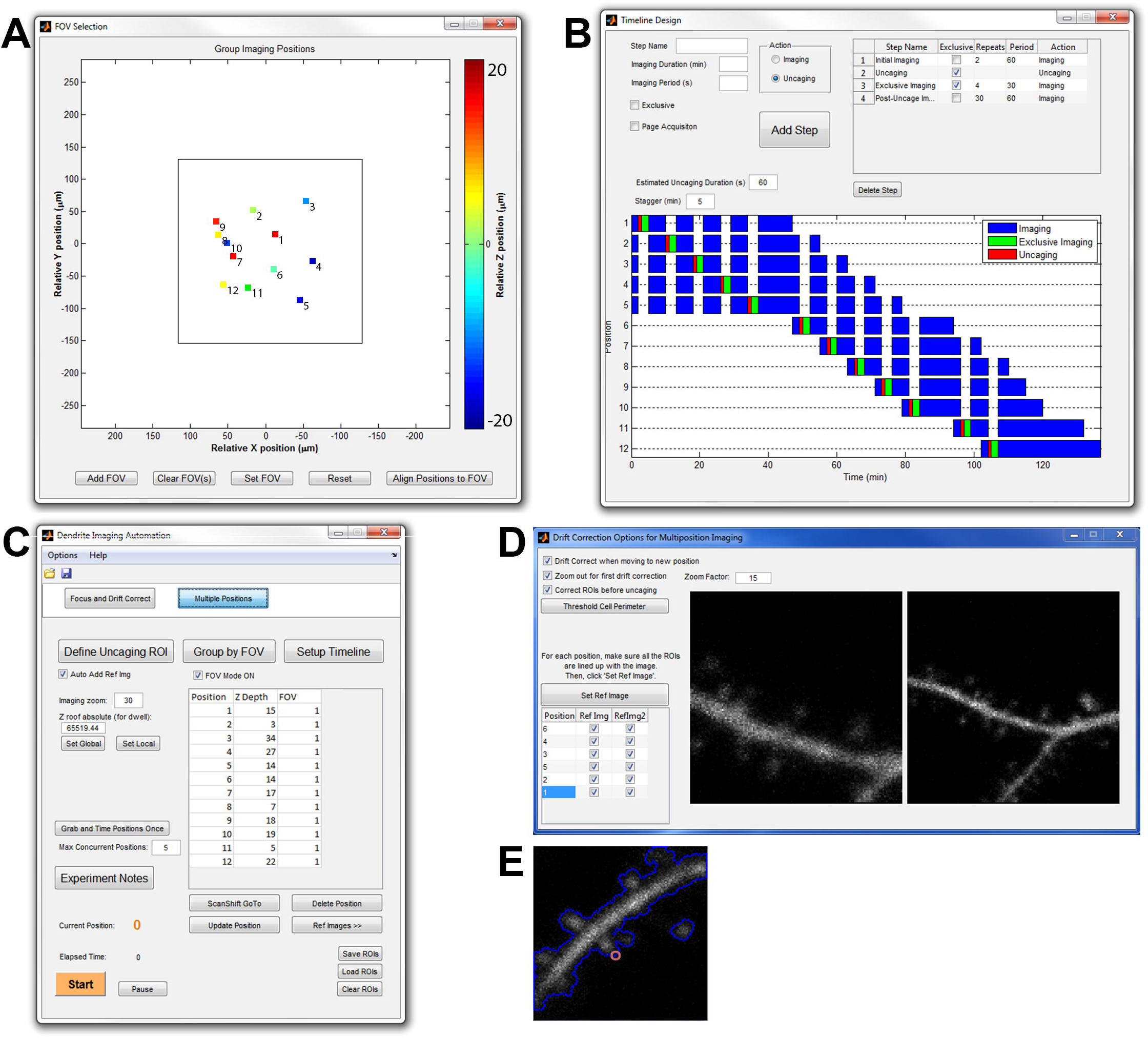
Non-motorized, automated, multi-position ROI selection, imaging, and photoactivation is controlled through a user-friendly interface. (A) GUI showing all motor positions that are translated to galvanometer scanning coordinates within a single field of view (FOV, large square). (B) A custom timeline interface allows users to design and preview imaging and (blue, green) and uncaging (red) cycles at each position. (C) A master GUI to keep track of and move between all imaging positions. Settings and coordinates can be saved and loaded. Z depth is set for each position to automatically modulate uncaging laser power to amounts of tissue interference in brain slices. Experimental Notes are automatically saved with each imaging cycle and can be altered to reflect experimental parameters. (D) GUI showing reference images for each position. Reference images are used for drift correction. Zoomed out reference images (right) are used for initial alignment. Threshold intensity values are set so each uncaging ROI (E, red) is shifted appropriately relative to the cell dendrite perimeter (E, blue).

To test and further optimize the efficacy of our non-motor multi-position imaging system, we measured structural plasticity of dendritic spines in hippocampal CA1 neurons transfected with enhanced green fluorescent protein (EGFP) in organotypic hippocampal slice prepared from mice (Fig 4). The lateral and axial drift corrections successfully tracked most spines and dendrites for a long time regardless of morphological changes. However, the spine set to be stimulated sometimes moves relative to the dendrite. Since glutamate uncaging had to be precisely targeted to the surface of the spine, we added a secondary drift correction method which would relocate the uncaging target to the surface of the spine immediately prior to uncaging. This allowed automated glutamate uncaging on multiple spines at consistent locations at the spine surface. We found that glutamate uncaging induced a rapid increase in volume within few minutes (transient phase) followed by long lasting enlargement (sustained phase), consistent with previous studies [3, 10, 41]. During the whole imaging session, the user did not need to be at the microscope, demonstrating the capability of automated tracking and stimulating software.

**Fig 4.**
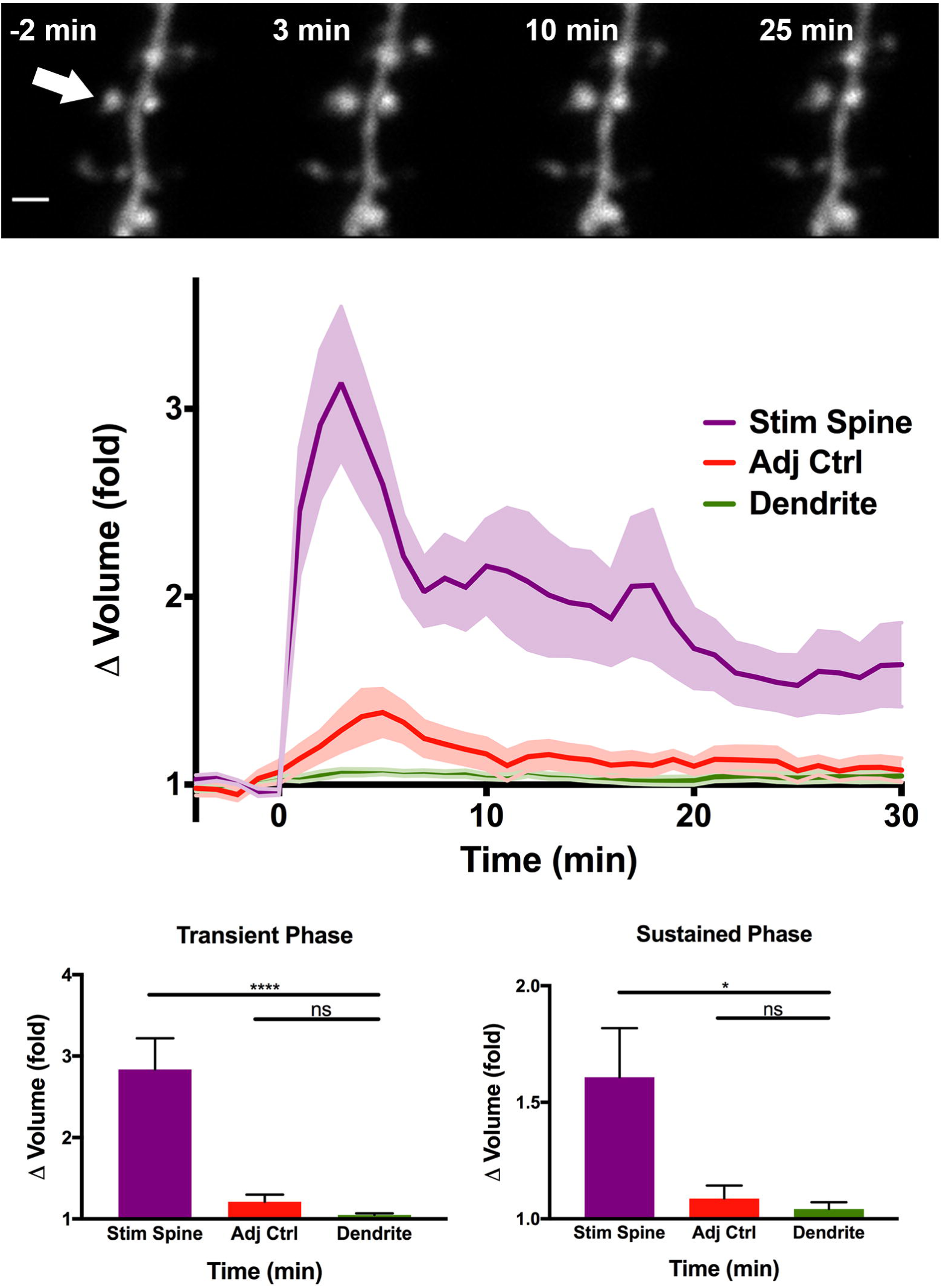
Plasticity in dendritic spines induced using automated focus, drift correction, and glutamate uncaging. (Top) CA1 dendrite pre- and post- uncaging. Arrow indicates photoactivation ROI. Scale = 1μm. Middle: Average volume change in spines following glutamate uncaging at t=0. Uncaging lasts 60s. Bottom: Quantification of transient (1–3 min) and sustained (26–30 min) change in spine volume. **** = p<0.0001, * = p<0.05. n=24 Stimulated spines, 7 neurons.

## Discussion

We have designed an easily implementable module for Scanimage to allow for multi-position scanning and photoactivation of dendritic spines to study postsynaptic plasticity. Furthermore, we described an implementation of an ETL in the excitation path which minimizes the signal loss and distortion. The ETL served both to increase imaging speed and remove sample drift caused by rapid stage or objective movements when changing focus. Our optical implementation of the ETL as a remote focusing element is combined with a straightforward user interface which is able to align and control the ETL current, and as a result, the axial focus. Furthermore, our software interface allows users to tilt the imaging plane in 3D by rapidly modulating the axial focus position in phase with the Y-scanning galvanometer. We expect this feature to be particularly useful in neuroscience, where long, straight neuronal projections can be scanned along their own axis, resulting in a significant increase in scanning efficiency. We found that an ETL-based system is relatively low price and easily implemented in any two-photon microscope. While the focusing speed of an ETL (15 ms) is sufficient for multi-position imaging, tilted-plane imaging needs to be performed fairly slowly, at around 1Hz, to allow the ETL sufficient refocusing time along the imaging plane.

It should be noted that there are faster (but more expensive and potentially more complicated) focusing devices which could be used with our software. For example, spatial light modulators (SLMs) can create 3D holograms within 2–4 ms, and have adaptive optics capabilities to potentially decrease distortion and aberration [42–44], A focusing device using a secondary objective and a galvanic mirror [45] provides even faster focus movement (∼ 1 ms). Perhaps the most well-established remote focusing element is the acoustic optic deflector (AOD), which can provide high-performance volume imaging in tissue [46, 47],

As for controlling software, we implemented a highly capable and customizable focus and drift correction system in order to broaden biological applications. Previously, dozens of autofocus algorithms have been made available through primary literature and open source code [20], However, these algorithms were not extensively implemented. Since imaging conditions may be drastically different for many users, we designed a tool capable of narrowing down the optimal algorithm based on accuracy and speed (Fig 3B). This allowed us to optimize our autofocusing algorithm to spine imaging experiments.

Finally, in order to allow the software to optimize experiments for biological events occurring at multiple time scales, we introduced a modular timeline scheduling feature which allows users to designate custom timeframes for imaging and photo-stimulation. This feature was especially important for studying sLTP, which has two distinct phases at different time scales (Fig. 4). During transient phase (first ∼5 min following stimulation), it is crucial to acquire images with higher frequency (typically ~10 − 60 s per image) since the volume change is rapid. However, during the steady state (~10 − 60 min), spine volume is stable and fast acquisition is unnecessary and rather damaging to the sample. The typical sampling time for the steady state is ~2 − 20 min. We demonstrated that this implementation allows us to measure the time course of sLTP in several spines (so far up to ~5 spines) with high efficiency.

## Conclusion

We successfully designed and implemented an automated system capable of reliably measuring sLTP in multiple dendritic spines. The implementation of an ETL in the excitation path, combined with galvanoic mirror scanning, allowed us to quickly switch between imaging positions with minimum perturbation to the sample. The customizable autofocus and drift correction system allows our software to track and stimulate individual dendritic spines over extended imaging sessions. By dramatically increasing the throughput of spine imaging and stimulation experiments, our system will accelerate studies to understand molecular basis of spine structural plasticity. In addition, the flexible implementation of software would allow researchers to use it for many imaging/photostimulation experiments.

## Methods

### ETL Implementation

An EL-10-30 ETL (Optotune) was implemented in the excitation pathway (Fig 1). The light path was designed in OpticStudio (Zemax) and optimized during setup so the ETL is conjugated to the back aperture of the objective. Lens L1 shapes the beam to fill the ETL. L2 and L3 are used to conjugate the ETL to the back aperture of the objective. A typical scan lens and tube lens setup passes the beam to the objective. The ETL is controlled by a current range of 0–300mA as indicated in the manual.

### Software Design

All programming was done in MATLAB to be compatible with Scanimage 3.8, available online for free (http://scanimage.vidriotechnologies.com). Autofocus and drift correction functions were implemented based on published code [29, 48], Abbreviations for focus measure operators are listed in Table 1. Maximum Brightness (BRGT) assigns focus values based on the maximum pixel intensity within the image. Maximum Brightness Gradient (MGRD) assigns focus values based on the maximum gradient magnitude value of the image. Drift correction in XY is calculated using a built-in MATLAB 2D cross-correlation algorithm [48],

### Evaluation of plasticity

Mouse pups were euthanized by deep anesthesia by isofluorane followed by decapitation. Organotypic hippocampal slice cultures were prepared as described previously [49] from p4-p6 mice and were cultured for 10–12 days before transfection. A biolostic particle delivery system (Helios^®^ Gene Gun System, Bio-Rad) was used to introduce fluorescent GFP labels to obtain sparse transfection of neurons. Two to six days after transfection, neurons in sparsely GFP-labeled CA1 hippocampal regions were chosen for imaging. Individual spines in the striatum radiatum on secondary apical dendrites were chosen for observation. MNI-caged L-glutamate (4-methoxy-7-nitroindolinyl-caged L-glutamate, Tocris) was uncaged with a train of 820-nm laser pulses (3.5–4 mW under the objective, 30 times at 1 Hz) near a spine of interest. Pulse duration was varied 4–8ms based on depth of the spine in tissue, allowing for reliable uncaging without excess light exposure. Experiments were performed at room temperature in ACSF solution containing (in mM): 127 NaCl, 2.5 KCl, 25 NaHC03, 1.25 NaH2P04, 4 CaCl2, 25 glucose, 0.001 tetrodotoxin (Tocris) and 4 MNI-caged L-glutamate, bubbled with 95% O2 and 5% CO2. All animal procedures were approved by the Max Planck Florida Institute for Neuroscience Animal Care and Use Committee, in accordance with guidelines by the US National Institutes of Health.

### Quantification and statistics

Spine volume was quantified using custom software written in MATLAB; all Z slices were summed together and oval and polygonal ROIs were drawn to select spines and dendrites, respectively. Volumes for each object were standardized to their average pre-uncaging values. Statistical significance was obtained using unpaired t-tests comparing the stimulated spine and adjacent spine averages to the dendrite as a control.

## References

1. Bailey CH, Kandel ER, Harris KM. Structural Components of Synaptic Plasticity and Memory Consolidation. Cold Spring Harb Perspect Biol. 2015;7(7):a021758. doi: 10.1101/cshperspect.a021758. PubMed PMID: 26134321.

2. Fu M, Yu X, Lu J, Zuo Y. Repetitive motor learning induces coordinated formation of clustered dendritic spines in vivo. Nature. 2012;483(7387):92–5. doi: 10.1038/nature10844. PubMed PMID: 22343892; PubMed Central PMCID: PMCPMC3292711.

3. Matsuzaki M, Honkura N, Ellis-Davies GC, Kasai H. Structural basis of long-term potentiation in single dendritic spines. Nature. 2004;429(6993):761–6. doi: 10.1038/nature02617. PubMed PMID: 15190253; PubMed Central PMCID: PMCPMC4158816.

4. Kim IH, Wang H, Soderling SH, Yasuda R. Loss of Cdc42 leads to defects in synaptic plasticity and remote memory recall. Elife. 2014;3:e02839–e. doi: 10.7554/eLife.02839. PubMed PMID: 25006034; PubMed Central PMCID: PMCPMC4115656.

5. Hayashi-Takagi A, Yagishita S, Nakamura M, Shirai F, Wu YI, Loshbaugh AL, et al. Labelling and optical erasure of synaptic memory traces in the motor cortex. Nature. 2015;525(7569):333–8. doi: 10.1038/nature15257. PubMed PMID: 26352471; PubMed Central PMCID: PMCPMC4634641.

6. Wilbrecht L, Holtmaat A, Wright N, Fox K, Svoboda K. Structural plasticity underlies experience-dependent functional plasticity of cortical circuits. J Neurosci. 2010;30(14):4927–32. doi: 10.1523/JNEUROSCI.6403-09.2010. PubMed PMID: 20371813; PubMed Central PMCID: PMCPMC2910869.

7. Colgan LA, Yasuda R. Plasticity of dendritic spines: subcompartmentalization of signaling. Annu Rev Physiol. 2014;76:365–85. doi: 10.1146/annurev-physiol-021113-170400. PubMed PMID: 24215443; PubMed Central PMCID: PMCPMC4142713.

8. Lee SJ, Escobedo-Lozoya Y, Szatmari EM, Yasuda R. Activation of CaMKII in single dendritic spines during long-term potentiation. Nature. 2009;458(7236):299–304. doi: 10.1038/nature07842. PubMed PMID: 19295602; PubMed Central PMCID: PMCPMC2719773.

9. Lisman J, Yasuda R, Raghavachari S. Mechanisms of CaMKII action in long-term potentiation. Nat Rev Neurosci. 2012;13(3):169–82. doi: 10.1038/nrn3192. PubMed PMID: 22334212; PubMed Central PMCID: PMCPMC4050655.

10. Murakoshi H, Wang H, Yasuda R. Local, persistent activation of Rho GTPases during plasticity of single dendritic spines. Nature. 2011;472(7341):100–4. doi: 10.1038/nature09823. PubMed PMID: 21423166; PubMed Central PMCID: PMCPMC3105377.

11. Rex CS, Chen LY, Sharma A, Liu J, Babayan AH, Gall CM, et al. Different Rho GTPase-dependent signaling pathways initiate sequential steps in the consolidation of long-term potentiation. J Cell Biol. 2009;186(1):85–97. doi: 10.1083/jcb.200901084. PubMed PMID: 19596849; PubMed Central PMCID: PMCPMC2712993.

12. Bayes A, van de Lagemaat LN, Collins MO, Croning MD, Whittle IR, Choudhary JS, et al. Characterization of the proteome, diseases and evolution of the human postsynaptic density. Nat Neurosci. 2011;14(1):19–21. doi: 10.1038/nn.2719. PubMed PMID: 21170055; PubMed Central PMCID: PMCPMC3040565.

13. Fernandez E, Collins MO, Uren RT, Kopanitsa MV, Komiyama NH, Croning MD, et al. Targeted tandem affinity purification of PSD-95 recovers core postsynaptic complexes and schizophrenia susceptibility proteins. Mol Syst Biol. 2009;5(269):269. doi: 10.1038/msb.2009.27. PubMed PMID: 19455133; PubMed Central PMCID: PMCPMC2694677.

14. Collins MO, Husi H, Yu L, Brandon JM, Anderson CN, Blackstock WP, et al. Molecular characterization and comparison of the components and multiprotein complexes in the postsynaptic proteome. J Neurochem. 2006;97 Suppl 1(m):16–23. doi: 10.1111/j.1471-4159.2005.03507.x. PubMed PMID: 16635246.

15. Redondo R, Bueno G, Valdiviezo JC, Nava R, Cristobal G, Deniz O, et al. Autofocus evaluation for brightfield microscopy pathology. J Biomed Opt. 2012;17(3):036008. doi: 10.1117/1.JBO.17.3.036008. PubMed PMID: 22502566.

16. Zhang ZC, Xia SR. [An autofocus algorithm based on principal component analysis]. Zhongguo Yi Liao Qi Xie Za Zhi. 2008;32(6):391–3, 7. PubMed PMID: 19253566.

17. Bravo-Zanoguera ME, Laris CA, Nguyen LK, Oliva M, Price JH. Dynamic autofocus for continuous-scanning time-delay-and-integration image acquisition in automated microscopy. J Biomed Opt. 2007;12(3):034011. doi: 10.1117/1.2743078. PubMed PMID: 17614719.

18. Shen F, Hodgson L, Hahn K. Digital autofocus methods for automated microscopy. Methods Enzymol. 2006;414:620–32. doi: 10.1016/S0076-6879(06)14032-X. PubMed PMID: 17110214.

19. Marks DL, Oldenburg AL, Reynolds JJ, Boppart SA. Autofocus algorithm for dispersion correction in optical coherence tomography. Appl Opt. 2003;42(16):3038–46. PubMed PMID: 12790455.

20. Geusebroek JM, Cornelissen F, Smeulders AW, Geerts H. Robust autofocusing in microscopy. Cytometry. 2000;39(1):1–9. PubMed PMID: 10655557.

21. Santos A, Ortiz de Solorzano C, Vaquero JJ, Pena JM, Malpica N, del Pozo F. Evaluation of autofocus functions in molecular cytogenetic analysis. J Microsc. 1997;188(Pt 3):264–72. PubMed PMID: 9450330.

22. Firestone L, Cook K, Culp K, Talsania N, Preston K, Jr. Comparison of autofocus methods for automated microscopy. Cytometry. 1991;12(3):195–206. Epub 01/01. doi: 10.1002/cyto.990120302. PubMed PMID: 2036914.

23. Price JH, Gough DA. Comparison of phase-contrast and fluorescence digital autofocus for scanning microscopy. Cytometry. 1994;16(4):283–97. doi: 10.1002/cyto.990160402. PubMed PMID: 7988291.

24. Fairley CR, Thompson TV, Lee KK. Method and apparatus for automatic focusing of a confocal laser microscope. Google Patents; 1997.

25. Jabbour JM, Malik BH, Olsovsky C, Cuenca R, Cheng S, Jo JA, et al. Optical axial scanning in confocal microscopy using an electrically tunable lens. Biomed Opt Express. 2014;5(2):645–52. doi: 10.1364/BOE.5.000645. PubMed PMID: 24575357; PubMed Central PMCID: PMCPMC3920893.

26. Chen JL, Pfaffli OA, Voigt FF, Margolis DJ, Helmchen F. Online correction of licking-induced brain motion during two-photon imaging with a tunable lens. J Physiol. 2013;591(19):4689–98. doi: 10.1113/jphysiol.2013.259804. PubMed PMID: 23940380; PubMed Central PMCID: PMCPMC3800448.

27. Grewe BF, Voigt FF, van't Hoff M, Helmchen F. Fast two-layer two-photon imaging of neuronal cell populations using an electrically tunable lens. Biomed Opt Express. 2011;2(7):2035–46. doi: 10.1364/BOE.2.002035. PubMed PMID: 21750778; PubMed Central PMCID: PMCPMC3130587.

28. Pologruto TA, Sabatini BL, Svoboda K. Scan Image: flexible software for operating laser scanning microscopes. Biomed Eng Online. 2003;2(1):13. doi: 10.1186/1475-925X-2-13. PubMed PMID: 12801419; PubMed Central PMCID: PMCPMC161784.

29. Pertuz S, Puig D, Garcia MA. Analysis of focus measure operators for shape-from-focus. Pattern Recognition. 2013;46(5):1415–32. doi: 10.1016/j.patcog.2012.11.011.

30. Shirvaikar M, editor An optimal measure for camera focus and exposure. Proceedings of the Thirty-Sixth Southeastern Symposium on System Theory (SSST04); 2004.

31. Harsh Nanda RC, editor Practical calibrations for a real-time digital omnidirectional camera. In Technical Sketches, Computer Vision and Pattern Recognition; 2001.

32. Helmli FS, Scherer S, editors. Adaptive shape from focus with an error estimation in light microscopy. Image and Signal Processing and Analysis, 2001 ISPA 2001 Proceedings of the 2nd International Symposium on; 2001: IEEE.

33. Lee S-Y, Yoo J-T, Kumar Y, Kim S-W. Reduced Energy-Ratio Measure for Robust Autofocusing in Digital Camera. IEEE Signal Processing Letters. 2009;16(2):133–6. doi: 10.1109/lsp.2008.2008938.

34. Krotkov E, Martin J-P, editors. Range from focus. Robotics and Automation Proceedings 1986 IEEE International Conference on; 1986: IEEE.

35. Pech-Pacheco JL, Cristóbal G, Chamorro-Martinez J, Fernández-Valdivia J, editors. Diatom autofocusing in brightfield microscopy: a comparative study. Pattern Recognition, 2000 Proceedings 15th International Conference on; 2000: IEEE.

36. Nikzad MSTSCA. Focusing Techniques. Journal of Optical Engineering. 1993:2824–36.

37. Eskicioglu AM, Fisher PS. Image quality measures and their performance. IEEE Transactions on communications. 1995;43(12):2959–65.

38. Nayar SK, Nakagawa Y, editors. Shape from focus: an effective approach for rough surfaces. Robotics and Automation, 1990 Proceedings, 1990 IEEE International Conference on; 1990: IEEE.

39. Thelen A, Frey S, Hirsch S, Hering P. Improvements in shape-from-focus for holographic reconstructions with regard to focus operators, neighborhood-size, and height value interpolation. IEEE Trans Image Process. 2009;18(1):151–7. Epub 12/20. doi: 10.1109/TIP.2008.2007049. PubMed PMID: 19095526.

40. Windsor RM, Univeresity of W, Abdul AM, Univeresity of W, Wu QMJ, Univeresity of W, et al., editors. 3D Shape from Focus and Depth Map Computation Using Steerable Filters. ICIAR; 2016: Springer Berlin Heidelberg.

41. Murakoshi H, Lee SJ, Yasuda R. Highly sensitive and quantitative FRET-FLIM imaging in single dendritic spines using improved non-radiative YFP. Brain Cell Biol. 2008;36(1–4):31–42. doi: 10.1007/s11068-008-9024-9. PubMed PMID: 18512154; PubMed Central PMCID: PMCPMC2673728.

42. Yang W, Miller JE, Carrillo-Reid L, Pnevmatikakis E, Paninski L, Yuste R, et al. Simultaneous Multi-plane Imaging of Neural Circuits. Neuron. 2016;89(2):269–84. doi: 10.1016/j.neuron.2015.12.012. PubMed PMID: 26774159; PubMed Central PMCID: PMCPMC4724224.

43. Lutz C, Otis TS, DeSars V, Charpak S, DiGregorio DA, Emiliani V. Holographic photolysis of caged neurotransmitters. Nat Methods. 2008;5(9):821–7. doi: 10.1038/nmeth.1241. PubMed PMID: 19160517; PubMed Central PMCID: PMCPMC2711023.

44. Nikolenko V, Watson BO, Araya R, Woodruff A, Peterka DS, Yuste R. SLM Microscopy: Scanless Two-Photon Imaging and Photostimulation with Spatial Light Modulators. Front Neural Circuits. 2008;2(December):5. doi: 10.3389/neuro.04.005.2008. PubMed PMID: 19129923; PubMed Central PMCID: PMCPMC2614319.

45. Botcherby EJ, Smith CW, Kohl MM, Debarre D, Booth MJ, Juskaitis R, et al. Aberration-free three-dimensional multiphoton imaging of neuronal activity at kHz rates. Proc Natl Acad Sci U S A. 2012;109(8):2919–24. doi: 10.1073/pnas.1111662109. PubMed PMID: 22315405; PubMed Central PMCID: PMCPMC3286923.

46. Duemani Reddy G, Kelleher K, Fink R, Saggau P. Three-dimensional random access multiphoton microscopy for functional imaging of neuronal activity. Nat Neurosci. 2008;11(6):713–20. doi: 10.1038/nn.2116. PubMed PMID: 18432198; PubMed Central PMCID: PMCPMC2747788.

47. Kirkby PA, Srinivas NKMN, Silver RA. Europe PMC Funders Group Europe PMC Funders Author Manuscripts A compact acousto-optic lens for 2D and 3D femtosecond based 2-photon microscopy. 2010;18(13):13721–45.

48. Sugar JD, Cummings AW, Jacobs BW, Robinson DB. A Free Matlab Script for Spatial Drift Correction. Microscopy Today. 2014;22(05):40–7. doi: 10.1017/s1551929514000790.

49. Stoppini L, Buchs PA, Muller D. A simple method for organotypic cultures of nervous tissue. J Neurosci Methods. 1991;37(2):173–82. PubMed PMID: 1715499.

